# STENCIL: A web templating engine for visualizing and sharing life science datasets

**DOI:** 10.1101/2021.06.04.447108

**Authors:** Qi Sun, Ali Nematbakhsh, Prashant K Kuntala, Gretta Kellogg, B. Franklin Pugh, William KM Lai

## Abstract

The ability to aggregate experimental data analysis and results into a concise and interpretable format is a key step in evaluating the success of an experiment. This critical step determines baselines for reproducibility and is a key requirement for data dissemination. However, in practice it can be difficult to consolidate data analyses that encapsulates the broad range of datatypes available in the life sciences. We present STENCIL, a web templating engine designed to organize, visualize, and enable the sharing of interactive data visualizations. STENCIL leverages a flexible web framework for creating templates to render highly customizable visual front ends. This flexibility enables researchers to render small or large sets of experimental outcomes, producing high-quality downloadable and editable figures that retain their original relationship to the source data. REST API based back ends provide programmatic data access and supports easy data sharing. STENCIL is a lightweight tool that can stream data from Galaxy, a popular bioinformatic analysis web platform. STENCIL has been used to support the analysis and dissemination of two large scale genomic projects containing the complete data analysis for over 2,400 distinct datasets. Code and implementation details are available on GitHub: https://github.com/CEGRcode/stencil

## Introduction

Advances in next-generation sequencing have supercharged biochemical assays into ‘big data’ genomic resources [1–3]. This explosion in data has been paralleled by the development of multiple quality control and data analysis tools [4–8]. The unique analysis requirements from each distinct genomic assay complicates the already diverse ecosystem of bioinformatic tools by necessitating the creation of a novel tools and algorithms to maximize biological interpretation [9–12]. Many of these tools are equipped with quantitative and qualitative metrics designed to analyze user-supplied data, generating insights into different aspects of the experiment. While many tools generate quality control (QC) reports, they do not always provide mechanisms for sharing these reports with the broader community or for generating reports composed of a diverse array of genomic experiments. Moreover, it is often not possible to programmatically access the curated data and visualized results that retain their original relationship to the source data

The Galaxy platform is a large-scale NSF and NIH-funded initiative intended to solve many of these issues [13]. Galaxy is a self-contained, portable, open-source software platform that creates sharable and executable script pipelines (workflows). Galaxy tracks the exact tool parameters and computational environment for every script and tool run on its platform to enable complete bioinformatic data reproducibility [14]. Novel bioinformatic tools are easily added in the Galaxy ecosystem with over 8,000 tools currently available (as of April 2021) for download and workflow execution [15]. Critically, Galaxy provides complete programmatic API control over the system for advanced users [16].

While the Galaxy platform has tremendous capabilities for running publication-validated workflows and generating reproducible data analysis, there is no current system in place to analyze the visualizations generated from multiple workflows simultaneously. The ability to readily inspect and interpret the data, without requiring programming expertise, plays a crucial role for advancing work in the life sciences, allowing wet bench scientists to validate experimental results, develop new hypotheses, and obtain insight into the massive output of genomic data provided by the latest technologies [17–20].

As a step toward data management and visualization, we developed STENCIL, a web templating engine that provides a framework for creating flexible, scalable, and interactive web visualizations that adapt to the growing needs of a project as required. It is designed for efficient reporting, data analysis visualization, and data dissemination in support of recommended FAIR Data practices [21]. STENCIL provides the ability to visualize and compare large sets of samples of interest within the same web frame. STENCIL supports retaining the links to reproducible workflows and removes any file storage redundancy, resulting in direct dataset visualization. It leverages Galaxy’s existing REST API interface to serve visualizations directly to STENCIL, providing complete programmatic control over all the displayed data, substantially improving a user’s discovery experience. Since STENCIL presents and aggregates the visualizations and analysis results already run from Galaxy or other workflow engines and pipelines, it is extremely light weight and can be run on an individual workstation or from a single CPU virtual machine.

## Design and implementation

### Software architecture

STENCIL is architected to aggregate data analyses from heterogeneous sources into a single reporting structure. STENCIL is composed of two primary subcomponents: Data Consuming Front End and Data Producing Back End. This design strategy dramatically simplifies horizontal scaling, allowing for the simultaneous running of multiple distinct STENCIL systems which can share the same core resources as specified. This high level of scalability ensures that STENCIL can serve thousands of unique datasets to multiple concurrent users on a simple webserver consisting of a single CPU and 8 Gb of RAM [22, 23]. STENCIL also supports common deployment strategies including Docker containers and locally managed servers through automated continuous integration/continuous deployment (CI/CD) workflows that are triggered by code changes.

STENCIL architecture is centered around ‘experiments’ as the fundamental unit. Each experiment contains one or more ‘sections’ that correspond to any arbitrary combination of data analysis output including, but not limited to: static images, interactive plots, and experimental meta-information in blocks of text or tables. Conceptually these sections consist of two elements: a React component forming the display element (frontend) and a JSON object serving as the data element (backend).

### Front end

STENCIL’s front end is a React JavaScript application that efficiently consumes REST APIs provided by the back end (**Figure 1A**). Importantly, there is no set requirement for visualization of any experiment. The STENCIL experiment page dynamically resizes itself to visualize the data that is attributed to each sample. As more data is added (i.e., over the course of a Galaxy workflow execution), static and interactive plots as well as data tables are added into an experiment’s front end in real-time.

**Figure 1.**
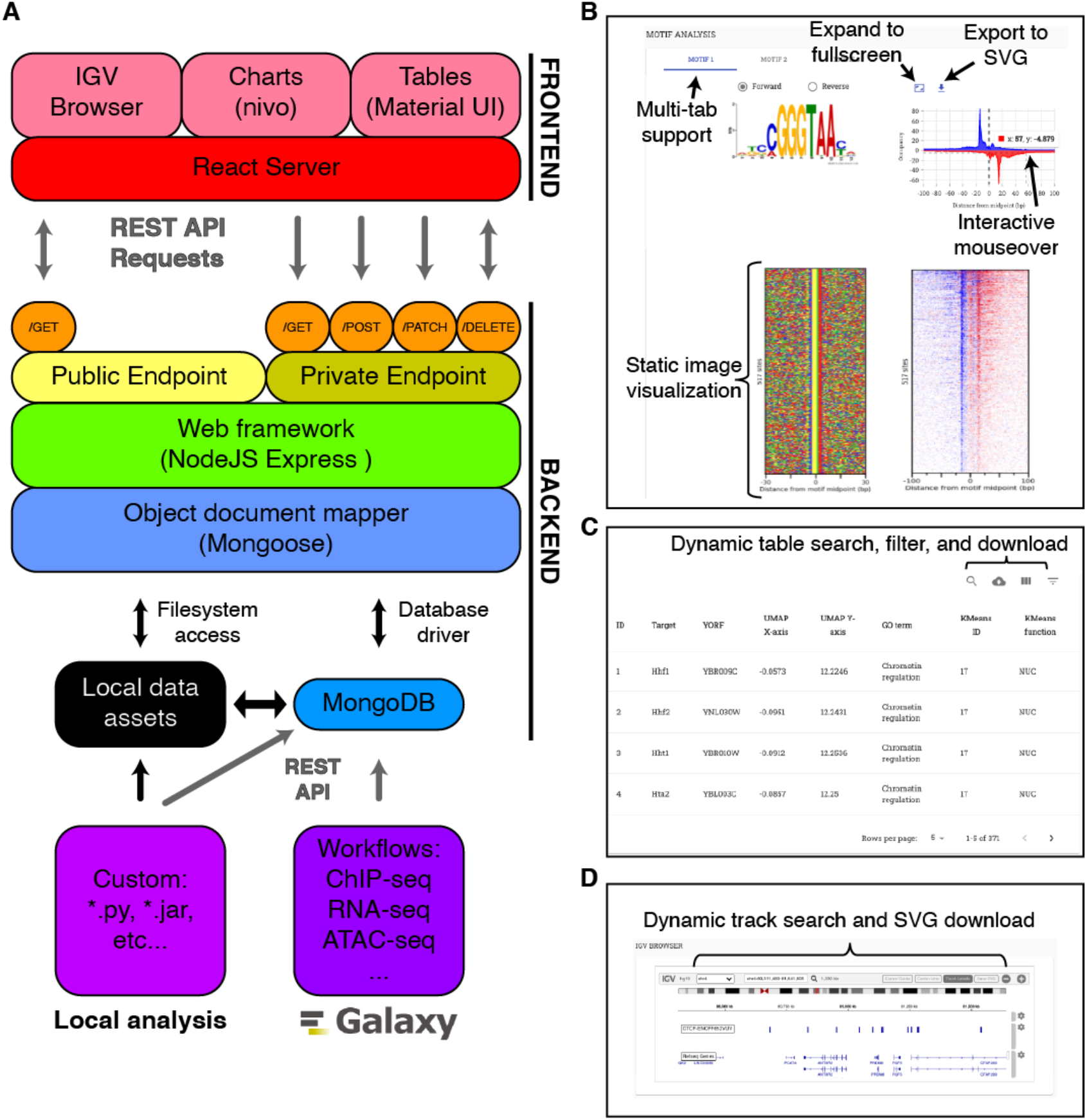
**A**, Overview of STENCIL architecture. A ReactJS frontend server provides web-access to the data. A NodeJS server and MongoDB instance store and manage the data provided and disseminated through RESTful API calls. Data analysis and hosted URLs are provided by the Galaxy platform and communicated directly to the MongoDB. Local analysis and file-hosting outside of the Galaxy platform is also supported. **B**, A sample React-served experiment section demonstrating multiple types of data. React serves static PNG/JPG/SVG images hosted remotely. Interactive charts include dynamic mouseover with additional data, the ability to expand to full screen, and export to SVG. **C**, Data table analysis allows for sorting and filtering using remotely hosted data. **D**, Integrated Genome Browser (igv) provides interactive track visualization in the same web frame as other advanced analysis.

While STENCIL provides a collection of default React components that effectively visualize common life science charts and images, extensibility was a primary consideration behind STENCIL’s design. As a result, the modular nature of STENCIL makes it is easy to rapidly add and remove new tool sections with minimal coding. The multiple plots of each sample can be organized into sections and tabs, with multiple sections per page, and multiple tabs per section. In each tab, the plots are organized either by rows or by columns. The number of rows and columns, as well as order of the plots in the layout can be customized in the configuration file using an intuitive syntax. Additionally, the prevalence of React in the scientific community, another key consideration in STENCIL’s design, directly translates to many existing React plotting tools that can be used for data visualization. The front end uses the d3.js-derived nivo charting library to create interactive dynamic charts from the data by default [24].

STENCIL comes with several pre-defined React components that can serve as initial templates for user-development. One of the sample templates is a React component that serves static PNG, JPG, and SVG images with an associated interactive JavaScript chart. Default chart features include dynamic mouseover with additional data, the ability to expand to full-screen, and an ‘export to SVG’ button for further refinement in vector-based imaging software (**Figure 1B**). Static images are generated outside of STENCIL by any number of pipelines and workflows including the Galaxy platform. The use of REST APIs provides complete programmatic control over all the displayed data. STENCIL supports basic access permissions to experiments and assigns each experiment a unique URL that can be publicly shared as needed.

Material UI-derived mui-datatables provide dynamic filtering and sorting of text-based data in table format [24] (**Figure 1C**). Genome browser functionality is provided through the integration of the BROAD Institute’s JavaScript implementation of IGV [25] (**Figure 1D**). Importantly, STENCIL is also ‘plug and play’ compatible with virtually all JavaScript charting libraries.

### Back end

STENCIL’s back end consists of a NodeJS Express application connected to a MongoDB database engine (**Figure 1A**). The MongoDB data models (**Figure 2A**) store and organize all the relevant meta-information associated with each experiment for display in the frontend, and an array of plot identifier and metadata associated with each plot. STENCIL’s backend database is designed to easily allow many-to-many relationships between experiments and projects, and between projects and users. The sample metadata include two required fields: project identifier and sample identifier; these are used to identify each experiment. The metadata for each plot include four required fields: layout identifier, tab identifier, plot identifier, and URL. The URL can be a link to the remote server or a local file path. The link can be either an image file or a JSON file with actual data values. If the link is a JSON file, the front-end plotting tool will be used to create a dynamic graph.

**Figure 2.**
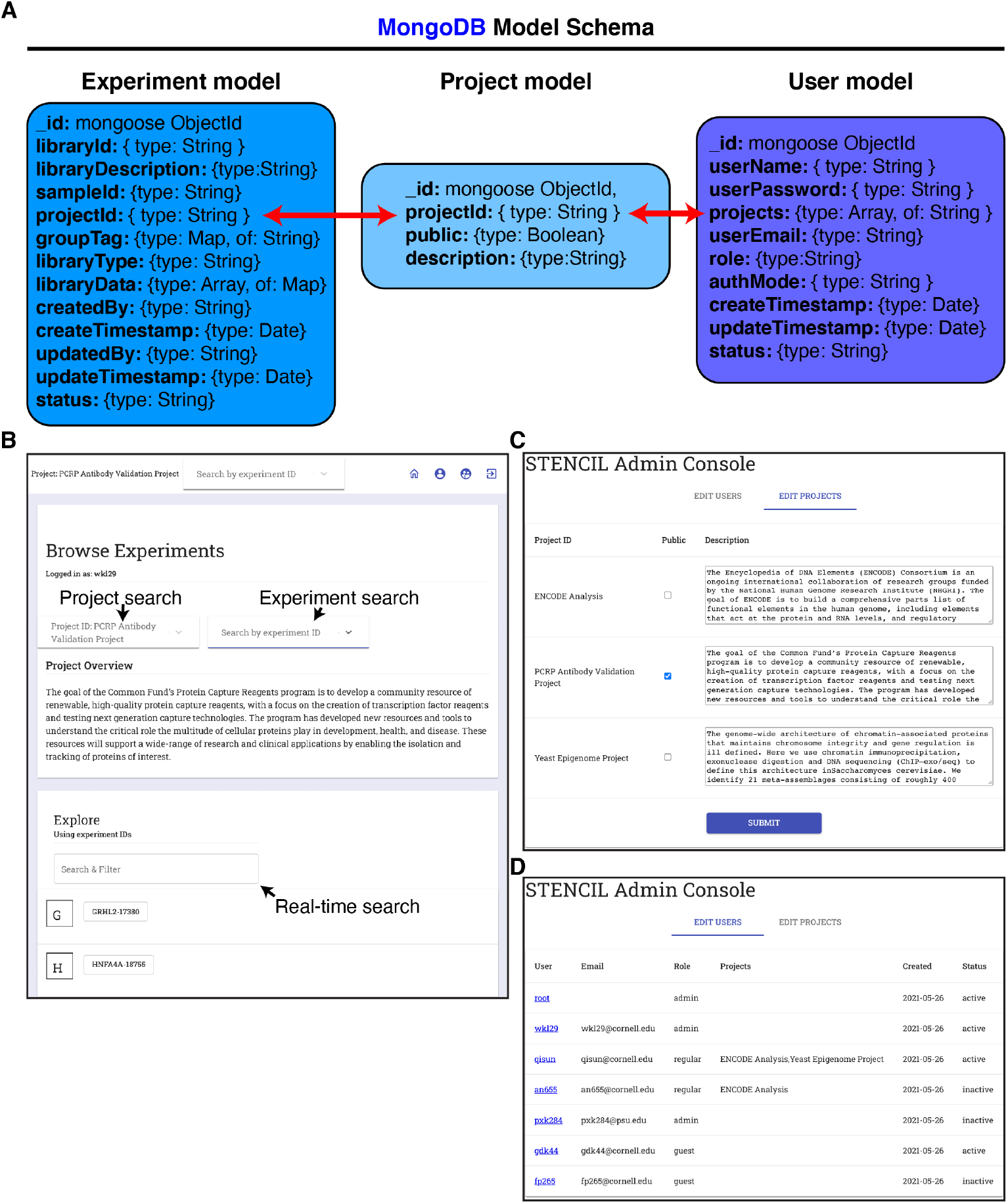
**A**, MongoDB model schema of STENCIL backend. URL references to experiment analysis are stored in the Experiment model. Experiments are linked to the Project model through a common projectId string. Users gain access to models and underlying experimental data through assignment to a variable number of projectIds. **B**, Experiment navigation screen of STENCIL. STENCIL supports users accessing multiple public and private projects as defined by the backend database. Real-time search enables quick filtering and access of experiments in large projects. **C**, STENCIL Admin console is available to STENCIL administrators and allows for project public/private control and project summary edits. **D**, STENCIL Admin console provides control of user access to projects and defining user roles.

STENCIL’s front end seamlessly parses these relationships and provides a user interface capable of navigating and modifying many of these relationships. The main landing page after login provides a user-searchable list of permission-assigned available projects (**Figure 2B**). Since each project can contain anywhere from a few dozen to a few thousand experiments [22, 23], the landing page also offers a real-time auto-complete searchable interface to find experiments within the project.

STENCIL provides Role Based Access Control to authorize appropriate access to the datasets and visualizations and provides more granular project-specific access control as well [26]. The application backend includes a permission authorization table in the MongoDB. Each ID is assigned a level of access to the application. Defined roles include Admin, Regular, and Guest.

A guest is a public user that may only view and download experimental data for projects that are designated as public. A Regular user is an authenticated user via a local login or Federated Single Sign-on. Regular users are only able to view and download public library pages until an administrator designates which project(s) they can work with on the website. Access to the Admin Console is granted to all designated STENCIL administrators and provides basic project and role capabilities. Administrators can make projects globally public, as well as edit the project summary displayed for each project (**Figure 2C**). The Admin role can define the level of access to a user ID; they can designate users to allow them to have appropriate access to view or edit the information. The Admin role also manages the projects, and assigns projects for each user, and can assign a Status to a User as ‘active’ or ‘inactive’ in order to more carefully control access to data files and experiment pages (**Figure 2D**).

### Galaxy integration

STENCIL was conceived of as a system to aggregate the downloaded results of multiple Galaxy workflows. This matured into its current iteration of hosting direct links to Galaxy datasets. In addition to the practical benefits of removing the need for data duplication (data already exists within Galaxy), it has the major benefit of maintaining the auditable and reproducible analysis generated within the Galaxy platform [14]. The datasets hosted on Galaxy enable STENCIL users to leverage Galaxy’s API and data dissemination features to achieve FAIR data standards compliance, all the way through final figure generation for publications.

Galaxy integration with STENCIL is enabled by two classes of custom Galaxy tools available here: (https://github.com/CEGRcode/galaxy_tools_for_stencil). The first class of tool is a pre-processing script that converts the output of an analysis tool into a JSON payload. Additional pre-processing scripts will be available to support common analysis algorithms (i.e., DESeq2, Cuffdiff, MEME). The second tool is a script whose sole purpose is to POST the plain-text JSON file to a STENCIL webserver (**Figure 3A**). The simplicity of the second tool provides enormous flexibility in its usage. The user pre-defines in a workflow (recommended), or using Galaxy’s real-time interface, whether the data being sent is a static image (PNG/JPG/SVG), a table, or a nivo chart to by dynamically rendered (**Figure 3B**). The current dynamic chart options supported are: Line Plot, Scatter Plot, Bar Plot, Heat Map, and Data Table and are selectable using a simple dropdown menu in the Galaxy interface. Additional chart options will be supported to accommodate a wider variety of supported analyses (https://usegalaxy.org/workflows/list_published).

**Figure 3.**
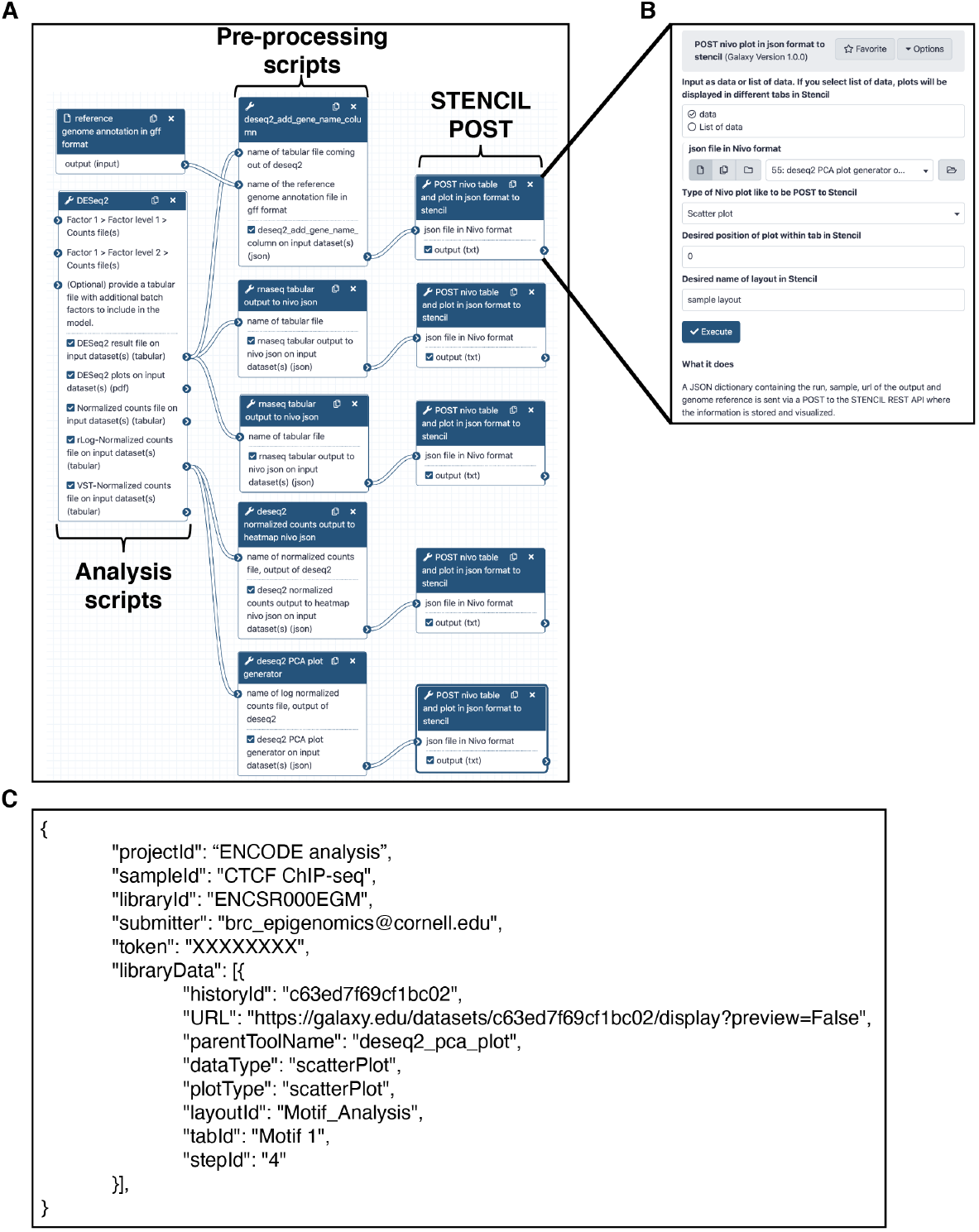
Galaxy integration with STENCIL **A**, A custom python script provides the mechanism through which a Galaxy tool (i.e., DESeq2) can POST its output to STENCIL **B**, The STENCIL-Galaxy communication tool allows the user to specify the type of data being transmitted to STENCIL **C**, A sample JSON payload containing the minimum amount of information needed by STENCIL to correctly place the analysis within an experiment and associated project.

The JSON payload sent from Galaxy contains the minimal amount of information required for STENCIL to properly visualize the data as well as provide a mechanism (Galaxy historyID) to trace the analysis back to Galaxy (**Figure 3C**). The small payload enables the STENCIL database to scale tremendously in size without dramatically increasing its disk space footprint. Additionally, the decision to use a document-based non-relational database (MongoDB) allows the user to trivially track any arbitrary additional sample meta-information such as genome ID, timestamp, workflow ID, etc. The design of this payloads allows for multiple Galaxy workflows to send their analysis results to the same STENCIL project and experiment. This structure fundamentally allows STENCIL to aggregate data from multiple discrete workflows into a single reporting structure.

### Alternative workflow support

Although STENCIL is primarily designed to complement existing Galaxy functionality, our architecture also supports integration with any tools capable of executing a POST request. This includes standard shell-script based analyses as well as alternative workflow engines such as Pegasus [27]. STENCIL is also able to seamlessly support file hosting from any arbitrary webserver (NGINX, Apache, etc.).

In the event a separate webserver is not available for hosting data, the included React backend server of STENCIL is also configured to allow for local file hosting. Critically, this functionality provides support for highly custom local analyses that often arise during algorithmic development and may not necessarily have existing support in the Galaxy toolshed. Although this process does not benefit from the significant bioinformatic tracking support provided by the Galaxy platform, it does allow bioinformaticians to rapidly iterate and develop novel algorithms, leveraging STENCIL’s visualization capabilities to develop robust and reproducible workflows.

### Data security and Single Sign-On integration

STENCIL provides two different authentication mechanisms: stand-alone user authentication and Shibboleth Single Sign-On (SSO). Upon installation of STENCIL, the default initial role of site administrator can set up either or both authentication options for the website. The application stores the login information in a cookie and allows for SSO login for an extended defined period.

For stand-alone user authentication, STENCIL stores the user ID and encrypted password within its backend MongoDB and authenticates the user by matching the stored password. The site administrator can reset local passwords and user IDs as needed in this model.

STENCIL can also be configured with the Internet2 standard Identity and Access Management system Shibboleth; Single Sign-On with integration to Two-Factor Authentication mechanisms. SSO provides the ability for users to authenticate to STENCIL using their university or institutional account, including integrated Two-Factor systems, without requiring additional management overhead for ID and password creation. This approach eases the cost of adoption of the software tool, particularly for smaller labs and individual PIs.

## Results

To date, STENCIL has been used to generate web applications to support analysis and dissemination of two large scale genomic projects each containing the data analysis for over 2,400 distinct genomic datasets. STENCIL currently operates on a webserver with a single CPU and 8Gb and easily hosts and disseminates all available data to multiple concurrent users.

### Protein Capture Reagents Program Validation

The Protein Capture Reagent Program (PCRP) was a NIH Common Fund project to generate and validate a renewable source of immunoreagents [28]. STENCIL was used to collate and provide a mechanism for the project’s data dissemination as per the NIH project requirements (http://www.pcrpvalidation.org) (**Figure 4**) [23]. Data was generated using a custom analysis pipeline (https://github.com/CEGRcode/PCRPpipeline) and uploaded to the STENCIL React backend. While this workflow enabled fast and accessible data analysis inspection, it lacked the full range of data reproducibility offered by the Galaxy platform.

**Figure 4.**
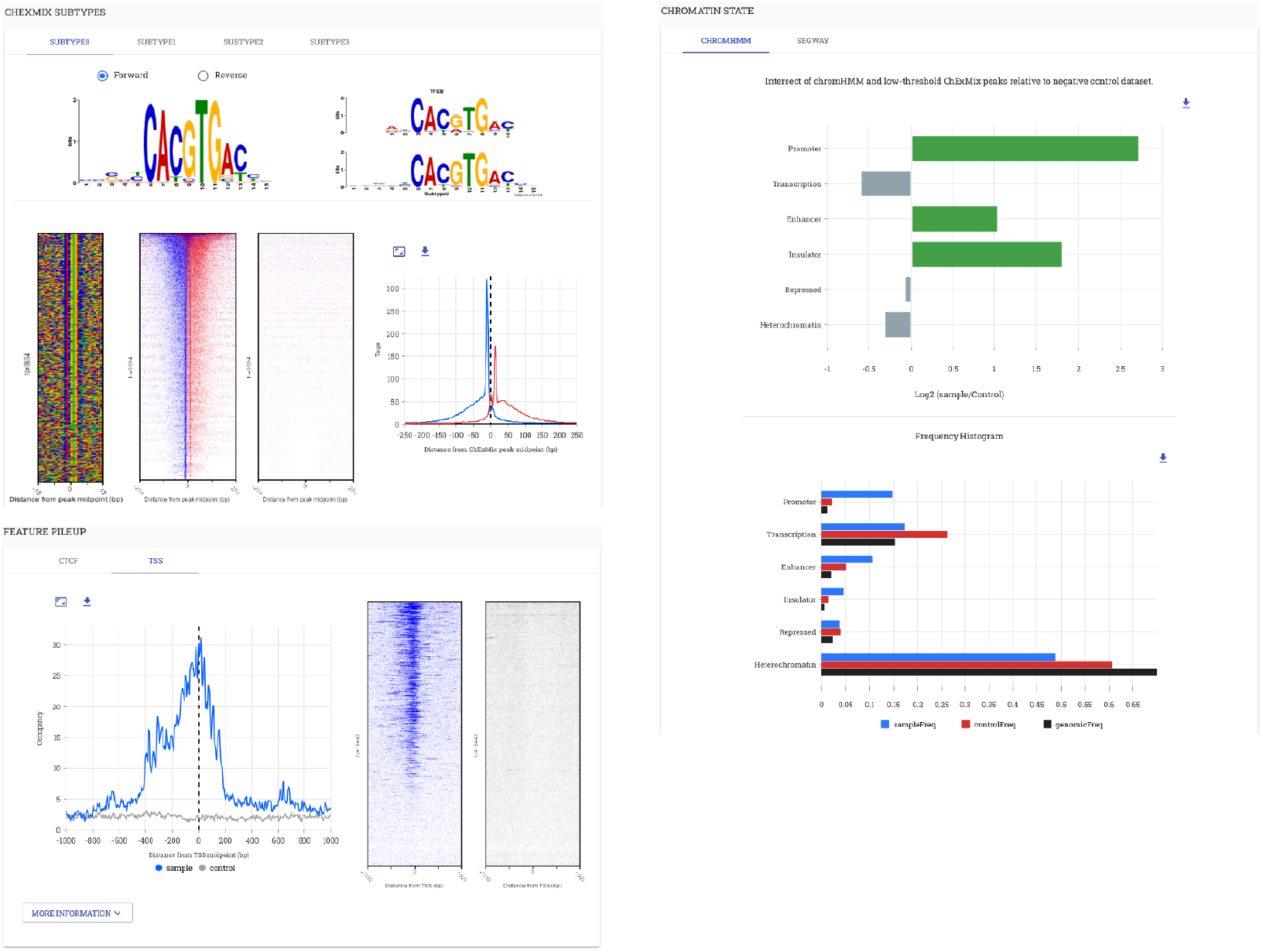
Application of STENCIL in the PCRP was an NIH-initiated project to screen 800 antibodies in ChIP-exo. NIH project requirements included providing the generated data in a community-accessible manner (http://www.pcrpvalidation.org).

### Yeast Epigenome Project

As part of our continued STENCIL development after the PCRP project, we incorporated Galaxy as a means of enabling completely reproducible data analysis. The Yeast Epigenome Project (http://www.yeastepigenome.org) was an ambitious endeavor to comprehensively map locations of protein-DNA interaction in *S. cerevisiae* [22]. All data for the Yeast Epigenome Project was generated using a Galaxy workflow available here: (https://github.com/CEGRcode/cegr-yep-qcviz/tree/master/exportedWorkflows). We generated over 1,200 unique genomic datasets and dozens of affiliated analyses per dataset that were all created using a completely reproducible Galaxy workflow. The resulting data was then uploaded to the React backend. STENCIL was used in this project to provide an interface for users to examine both quality control metrics as well as interrogate the baseline biology of each unique factor’s binding in the genome (**Figure 5**).

**Figure 5.**
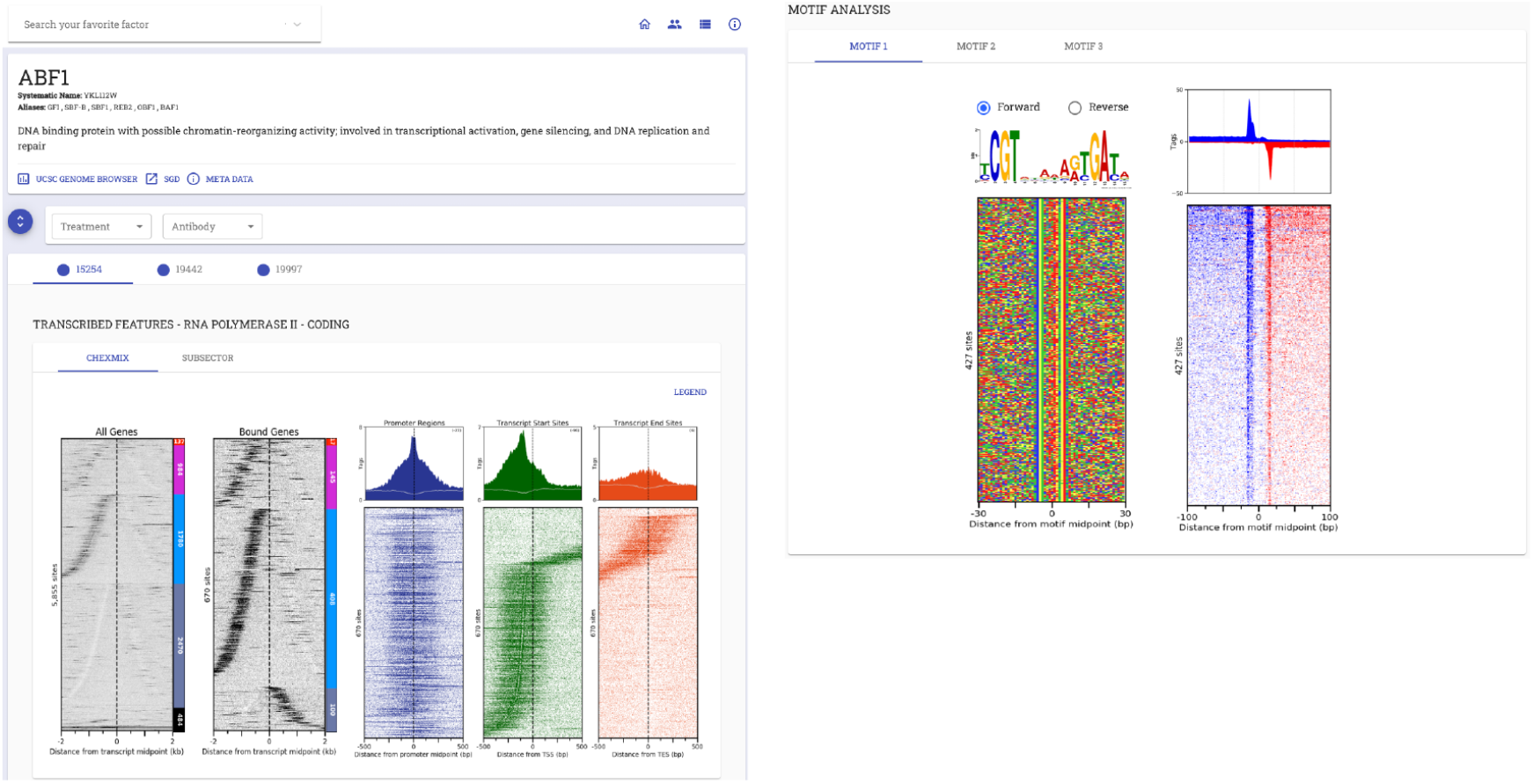
Application of STENCIL in the Yeast Epigenome Project. A comprehensive map of protein binding in *S. cerevisiae* required the development of a Galaxy workflow to both generate quality control metrics as well as provide baseline biological insight. Full analysis with high-resolution figures is available at http://www.yeastepigenome.org/yep/factor/ABF1.

### Results of RNA-seq and ATAC-seq pipelines in STENCIL

We continued to refine our use of STENCIL to remove data duplication resulting from downloading data from Galaxy and uploading to STENCIL, although we maintain that functionality to support all possible analysis workflows. The recommended usage of STENCIL is to host data directly from Galaxy. We provide a template RNA-seq analysis workflow that POSTs the data URL directly from Galaxy instance to a STENCIL webserver. Since the data is stored and hosted from Galaxy, there is no unnecessary data duplication. As an additional advantage, in this configuration, STENCIL now maintains a link to the unique Galaxy history and data. This provides a continuous link from initial data input all the way through figure generation in line with FAIR data practices. In addition to being able to host thousands of concurrent datasets, STENCIL can visualize tens of thousands of datapoint simultaneously and interactively in the same web frame (**Figure 6B**). We have also the applied the same workflow using ATAC-seq data demonstrating its utility across multiple genomic assays (**Supp Figure 1**).

**Figure 6.**
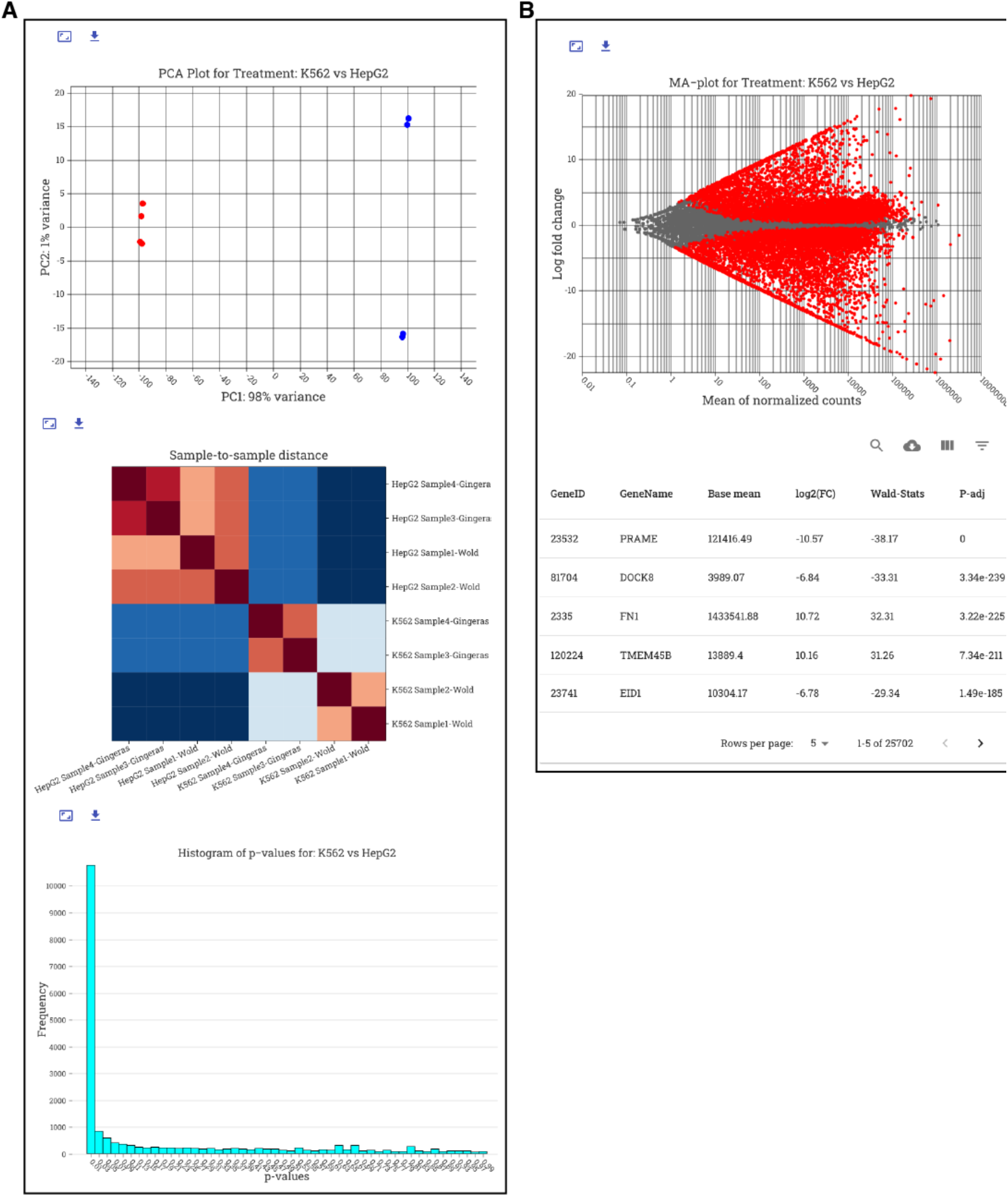
DESeq2 RNA-seq analysis visualized in STENCIL. **A**, DESeq2 charts are visualized as dynamic charts in STENCIL. All data is generated and hosted directly from Galaxy. **B**, The nivo charting library allows for on-the-fly generated of interactive plots containing tens of thousands of unique datapoint in seconds.

## Discussion

One of the main bottlenecks in analysis of genomic data is efficient and scalable visualization approaches. To address this challenge, we developed a graphical reporting interface (STENCIL) designed to integrate data generated by the Galaxy platform or in combination with custom lab workflows. STENCIL is capable of dynamically composing interactive graphical reports on biological results from multiple genomic assays. In its Galaxy integration option, STENCIL displays its data directly from the Galaxy instance, removing the need for local duplication of data files. STENCIL also enables aggregate data from multiple assays and workflows from any hosted Galaxy instance into a single reporting structure. The data generated by Galaxy is visualized interactively and provides capability for user-downloads of pre-packaged datasets and publication-quality figures for further analysis. Users can alternatively download STENCIL-analyzed data in a format compatible with analysis programs such as PRISM and Microsoft Excel.

Future development of STENCIL, in addition to generating automated reports for high-throughput experimental pipelines, will provide additional options for users to interactively work with the data to create customized reports based on user-selected samples and user-selected options for further analyses without having to leave the front-end web browser environment. A simple webform will enable users to select datasets of interest, or to upload other datasets, then select compatible analysis workflows. Analyses will then be performed on remote HPC systems to generate and subsequently visualize the data and results on the fly [29]. STENCIL will be further expanded to include quality control metrics and biological discovery workflows associated others popular genomic experiments (i.e., scRNA-seq, GWAS). The authors welcome and encourage support and contributions by the larger life sciences community, particularly the involvement of the active Galaxy.org open-source community. Future development includes plans for sharing out this tool and conducting training to the Galaxy and other HPC research communities.

STENCIL is designed to efficiently report data analysis, generate high-quality visualizations for figures, and disseminate the corresponding data while ensuring FAIR Data practices [21]. The ability to visualize and compare samples of interest from multiple distinct workflows within the same web frame assists researchers’ interpretation of results from inter-relating large-scale datasets. This coalescence of analysis should result in better evaluation of signal-versus-noise (experimental quality control) when visualizing the results. The multiple mechanisms that STENCIL provide for reporting and visualization will facilitate sharing and utilization of reproducible life science data. Reports produced by STENCIL should allow the researchers to gauge the underlying biology from the experiment in an efficient and scalable manner.

---

**Supp Figure 1.**
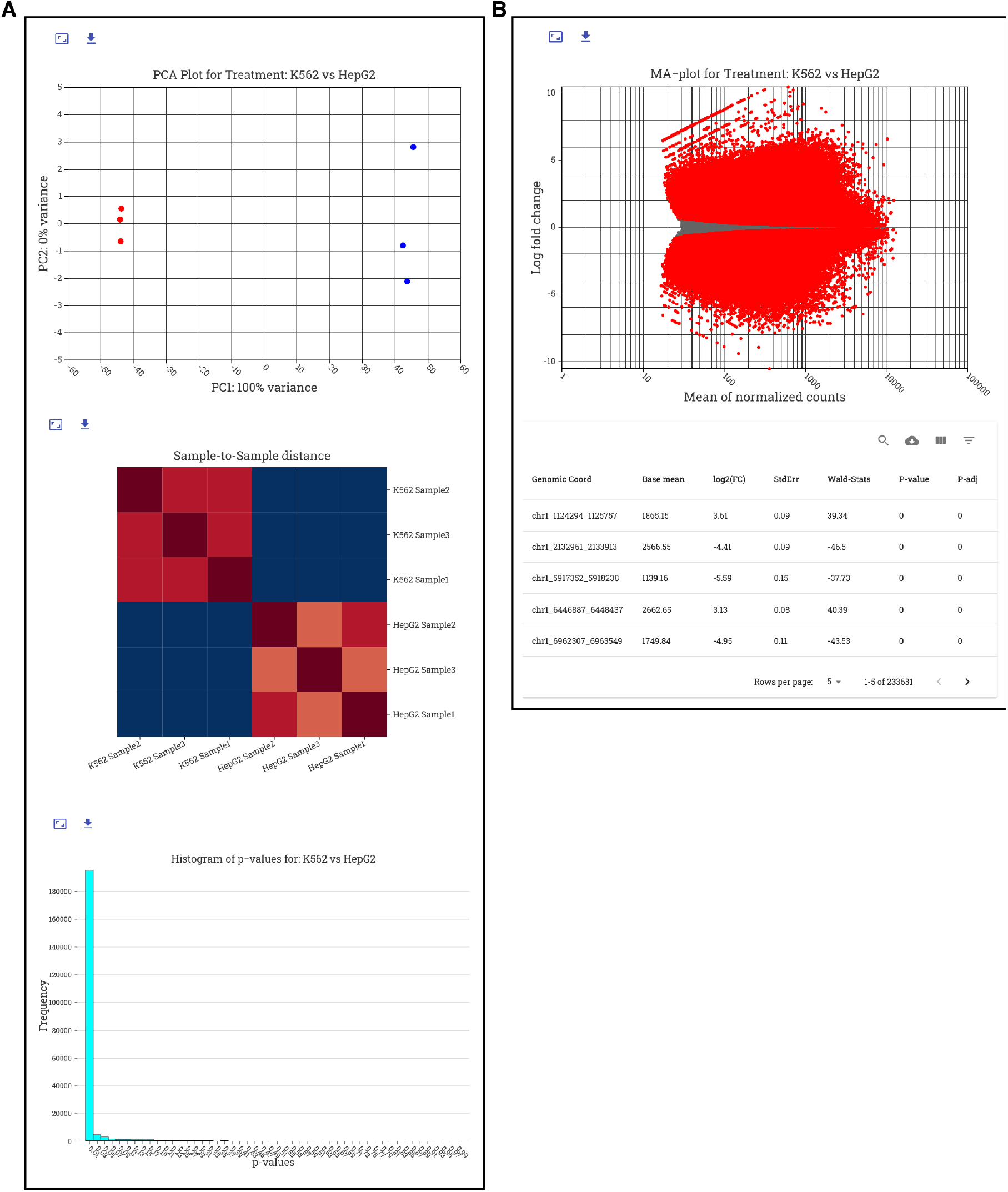
DESeq2 ATAC-seq analysis visualized in STENCIL. **A**, DESeq2 charts are visualized as dynamic charts in STENCIL. All data is generated and hosted directly from Galaxy. **B**, The nivo charting library allows for on-the-fly generated of interactive plots containing hundreds of thousands of unique datapoint in seconds.

## Supporting information

Source code for STENCIL is available at https://github.com/CEGRcode/stencil with detailed documentation and examples are available at https://CEGRcode.github.io/stencil/. Sample Galaxy tools to communicate with STENCIL are available at https://github.com/CEGRcode/galaxy_tools_for_stencil.

## Acknowledgements

The work was supported through the Cornell Institute of Biotechnology’s Epigenomic Core Facility, (https://www.biotech.cornell.edu/core-facilities-brc/facilities/epigenomics-facility).

